# Role of TNF-α among inflammatory molecules secreted by injured astrocytes in the modulation of *in vitro* neuronal networks

**DOI:** 10.1101/2020.10.16.341644

**Authors:** Joséphine Lantoine, Anthony Procès, Agnès Villers, Sophie Halliez, Luc Buée, Laurence Ris, Sylvain Gabriele

## Abstract

Traumatic brain injury (TBI) remains one of the leading causes of mortality and morbidity worldwide. Despite its high prevalence and extensive efforts to develop neuroprotective therapies, effective treatments for TBI are still limited. Among important neuronal damages, TBI induces structural and functional alterations of astrocytes, the most abundant cell type in the brain. Injured astrocytes respond in diverse mechanisms that result in reactive astrogliosis and are involved in the physiopathological mechanisms of TBI in an extensive and sophisticated manner. The establishment of effective neuroprotective treatments for TBI requires to better understand the complex biochemical interactions between activated astrocytes and neurons that contribute to the secondary injury. To address this challenge, we studied *in vitro* the role of mechanically injured astrocytes on the growth and synaptic connections of cortical neuronal networks of controlled architectures grown on well-defined protein micropatterns. Astrocytes were cultivated on elastic membranes and mechanically activated by stretching cycles. The culture media of healthy or activated astrocytes was then introduced on neuronal networks. We analyzed the neuronal viability, the neurite growth and the synaptic density of neuronal networks to understand the role of the inflammatory molecules secreted by mechanically activated astrocytes. Furthermore, we cultivated neuronal networks during 13 days with different doses of TNF-α in order to decipher its individual contribution among the other cytokines. Here we show that the ratio of tubulin to synapsin area was significantly higher in neuronal networks treated with either 4 or 2 doses of TNF-α, suggesting that TNF-α can promote the tubulin polymerization process. Assuming that TNF-α can bind to either TNFR1 or TNFR2 receptors, which lead respectively to the cell survival or the cell apoptosis, we studied the modulation of the both TNF-α receptors in response to the medium of mechanically activated astrocytes and different doses of TNF-α. Our findings indicate that the amount of both receptors increases with the maturation of the network. In addition, we observed a significant modulation of the amount of TNFR1 and TNFR2 in response to the media of injured astrocytes that leads to a large imbalance between both receptors, suggesting an important role for TNFα-signaling in the physiopathological mechanisms of TBI.

## 1 Introduction

It is now well established that a mechanical stress applied to the central nervous system (CNS) induces glial activation and neuroinflammation processes leading to a cascade of complex and interdependent events. These events give rise to the clinical manifestations of the traumatic brain injury (TBI) (Moshayedi, 2014; Hemphil, 2015). The initial inflammatory process induced by TBI has been shown to progress to chronic traumatic encephalopathy (Perrine, 2017; Hay, 2016) which can persist on a long period of time till several years in some clinical cases (Gentleman, 2004; Johnson, 2013; Ramlackhansingh, 2011; Chen, 2009; Hayes, 2017). However, the neuroinflammatory response to TBI possesses both beneficial and detrimental effects, and these are likely to differ in the acute and delayed phases after injury (Smith, 2013). After having proliferated at the injury site, microglial cells along with astrocytes fuse to form an area of containment between healthy and injured tissues called the glial scar (Sofroniew, 2009; Puntambekar, 2018). Microglia and astrocytes switch to a reactive state and play their role in both neuroinflammation and tissue healing processes. When microglia become reactive, it can induce detrimental neurotoxic effects by releasing multiple cytotoxic substances, including proinflammatory cytokines, such as interleukin-1b [IL-1b], tumor necrosis factor-α [TNF-α] or interferon-c [IFNc] (Block, 2007).

Besides, activated microglia is also involved in neuronal regeneration but only upon activity regulation. This regulation is done, among others by the astrocytes and the soluble factors released in the extracellular compartment. The release of proinflammatory cytokines and other soluble factors by activated microglia may significantly influence the subsequent activation of astrocytes and glial scar formation. Evidence points therefore towards an important role of astrocytes in traumatic brain injury, both at an early stage and at later stages during glial scar formation (Burda et al. 2016; Karve et al. 2016). Indeed, the presence of mechano-sensitive channels in the membrane of astrocytes induced an immediate response of these cells to mechanical injury (Islas et al. 1993). Reactive astrocytes release ATP which in turn can activate intracellular pathway in glial cells, neurons and vascular cells (Neary et al. 2005; Verderio et al. 2001). On the other hand, it has also been shown that intermediate filament, such as vimentin and GFAP can drive astrocyte reactivity and beneficial effect on axonal repair after traumatic brain injury (Liu et al 2014; Pekny et al. 1999).

Depending on their state of activation and on the cellular microenvironment, reactive astrocytes, as microglial cells, can release various inflammatory molecules, which could have a detrimental or beneficial effect on neural networks (Burda et al. 2014). The role of astrocytes in neuronal network development is clearly demonstrated (Clarke et al 2013) and Tyzack et al. have shown their role in synaptic plasticity following motor nerve injury (Tyzack etal. 2014). More precisely, TNF-α released by astrocytes in control conditions (Beattie et al 2002) or after denervation (Becker et al. 2013) was showed to maintain homeostatic synaptic plasticity in neuronal networks. At later stages, reactive astrocytes play an important role in the regulation of immune cells infiltration, blood-brain barrier integrity, glial scar formation and cerebral blood flow regulation (Bush et al. 1999; Abbott et al. 2006; Villapol et al 2014). Moreover, astrocytes have an important role in the regulation of extracellular glutamate preventing excitotoxicity induced by neuronal damage (Chen et al. 2003) and release neurotrophic factors preventing neurodegeneration after brain injury (Madathil et al. 2013; Myer et al 2006). However, other studies underlie the detrimental effect of astrocytes and glial scar on neuronal survival after fluid percussion injury (Di Giovanni et al. 2005)

As the traumatic brain injury is one of the leading cause of mortality and morbidity with 10 million of death annually (Langlois, 2006) and with 5.3 million people living with a TBI-related disability (Simon, 2017), it is therefore important to cover the lack of efficacious treatments focused on the neurotoxic pathways that occurs after the primary injury leading to an aggravation of the injury and to long-term impact. The key to develop future neuroprotective treatments is then to target post-traumatic neuroinflammation and microglial activation by minimizing the detrimental and neurotoxic effects of neuroinflammation, while promoting beneficial and neurotrophic ones. In order to create optimal conditions for regeneration and repair after injury, it is essential to better understand the molecular events that contribute to secondary injury and to study the effect of inflammatory processes on the functioning of the neural network. The complexity of the *in vivo* situation prevents from individually manipulating important parameters of the neuroinflammation process that occurs in response to a brain injury and often leads to limited access to the cellular tissue of interest. The study of the inflammation response in TBI requires standardized *in vitro* cellular models and mechanical injury conditions, respectively.

In this study, we studied *in vitro* the role of mechanically injured astrocytes on the growth and synaptic connections of neuronal networks. To address this challenge, we used cortical neuronal networks of controlled architectures grown on well-defined protein micropatterns (Lantoine et al. 2016). In addition, astrocytes were cultivated on stretchable devices and their culture media (healthy or mechanically activated) were introduced on healthy developing neuronal networks. Then we used an electrochemiluminescence method to determine the concentrations of the different cytokines in culture media of the mechanically-activated astrocytes. By staining GFAP in astrocyte cultures, we showed that two consecutive stretch cycles can mechanically activate astrocytes. We analyzed the neuronal viability, the neurite growth and the synaptic density of developing neuronal networks to understand the role of the inflammatory molecules secreted by mechanically-activated astrocytes. Then, we cultivated neuronal networks during 13 days with different doses of TNF-α in order to decipher the individual contribution of TNF-α. Our findings indicated that TNF-α do not trigger cell death, whereas that the ratio of the tubulin to the synapsin area was significantly higher in neuronal networks treated with either 4 or 2 doses of TNF-α, suggesting that TNF-α can promote the tubulin polymerization process. Assuming that TNF-α can bind to either TNFR1 or TNFR2 receptors, which lead respectively to the cell survival or the cell apoptosis, we studied the modulation of the both TNF-α receptors in response to the medium of mechanically-activated astrocytes and different doses of TNF-α.

## 2 Material and Methods

### 2.1 Preparation of cell culture substrate

Cellular substrates of 500 kPa in Young’s modulus were made in polydimethylsiloxane (PDMS), which has increasingly been employed for the fabrication of neuronal cell culture platforms and microfluidic devices (Grevesse et al. 2015; Lantoine et al. 2016). PDMS curing agent (Dow Corning, Sylgard 184) was mixed with a base agent in a mass ratio of 22:1 in 15 ml centrifugal tubes. The mixture was placed in a vacuum for 30 min to remove air bubbles and a volume of 100 μl was transferred using a micropipette on the top of a 25 mm glass coverslip. Then the PDMS mixture was spin-coated with a speed gradually increasing from 500 to 5000 rpm, for a total duration of 1 min and cured at 60°celsius for 4 hours. The obtained samples exhibit a thickness of 25 μm (Fig. 1A). Samples were then stored at room temperature in a vacuum desiccator (Versaevel, 2014). The Young’s modulus of the PDMS elastomers were measured by DMA (Dynamic Mechanical Analysis, Mettler Toledo DMA/SDTA 861e, Switzerland) in compression mode on circular cylindrical samples of 15 mm in diameter and height of 10 mm. Samples were sandwiched between two parallel plates and an oscillating strain of maximum amplitude of 10 % was applied and the stress needed to deform the cylindrical samples was measured over a frequency range of 0.1-10 Hz. During compression testing, a settling time of approximately one minute was used to achieve a stable measurement of the storage modulus at each frequency. We determined a Young’s modulus of 508 ±16 kPa (n=13).

**Figure 1.**
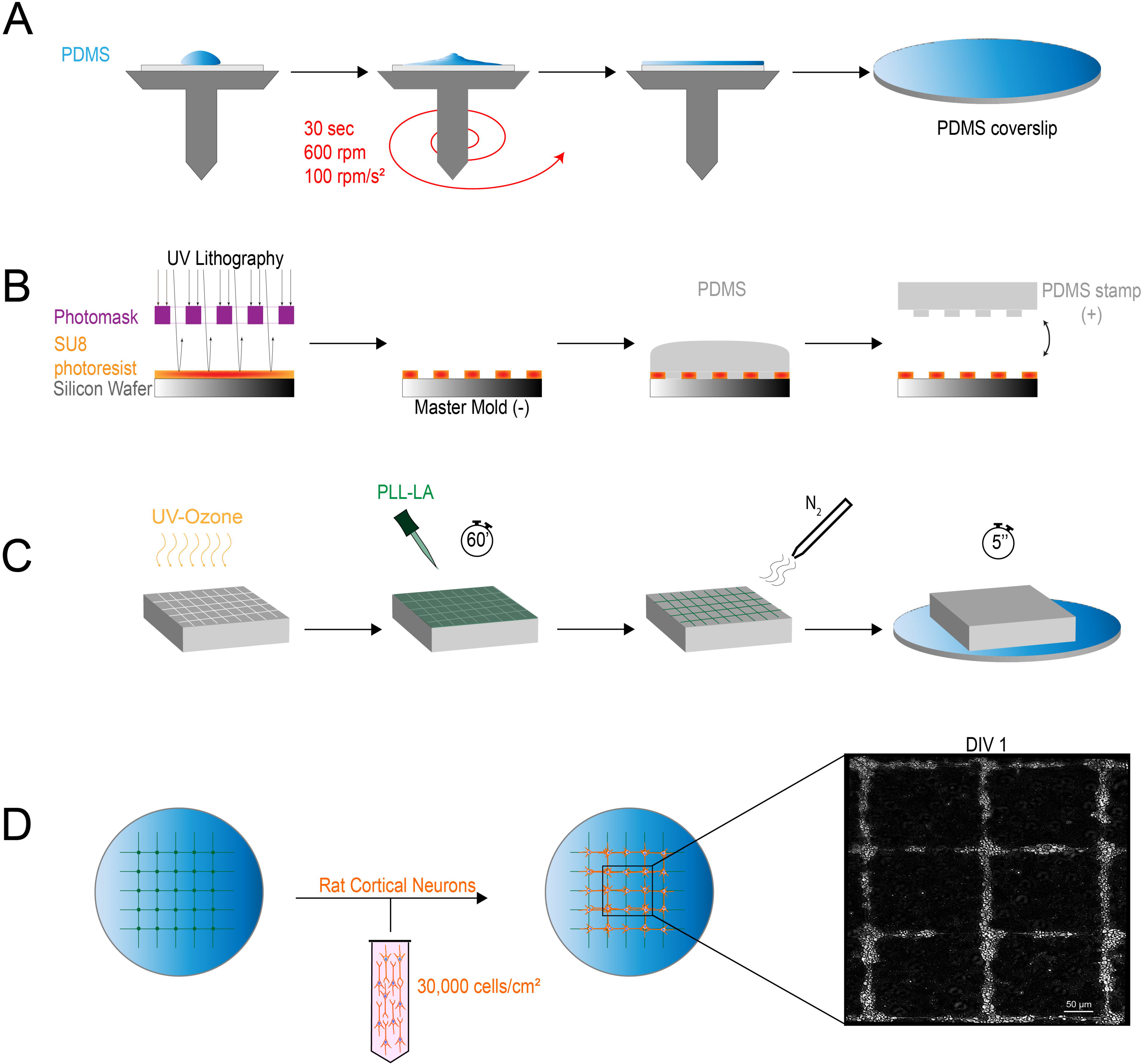
Neuronal networks of primary cortical neurons were culture on micropatterned substrates. (A) Polydimethylsiloxane (PDMS) is dropped on a glass coverslip and spin-coated to obtain a 25 μm thick membrane. (B) Soft lithography techniques were used to make PDMS microstamps. (C) PDMS stamps were incubated with a solution of poly-L-lysine (PLL) and laminin (LA) and dried with nitrogen. PDMS-coated glass coverslips were stamped to create a PLL/LA micropattern. Non-printed regions were passivated with Pluronics to avoid cell adhesion. (D) Rat cortical neurons (RCN) were seeded on the microprinted coverslip and allowed to grow at least 24 hours before changing medium.

### 2.2 Microcontact printing

PDMS substrates were microprinted with a 50:50 solution of poly-L-lysine (PLL, P4707, Sigma) and laminin (LA, L2020, Sigma). Flat PDMS microstamps were prepared by casting a 10:1 (w/w) degassed mixture of PDMS prepolymer and curing agent (Sylgard 184, Dow Corning) on an engraved silicon master, which was passivated with fluorosilane (tridecafluoro-1,1,2,2-tetrahydrooctyl-1-trichlorosilane, Gelest) in a vacuum to facilitate the removal of the PDMS layer (Figure 1B). After curing overnight at 65°C, flat PDMS microstamps were peeled off the silicon wafer, washed with a detergent (5 % Decon) then rinsed with distilled water and finally washed with 70 % ethanol to remove the remaining silane. To made them hydrophilic, microstamps were exposed during 7 min to ultraviolet/ozone (UV/O_3_). Activated PDMS microstamps were then incubated with a sterile solution of PLL-LA for 1 hour at room temperature and then dried with pure nitrogen (Figure 1C). Inked PDMS microstamps were then gently applied on flat PDMS surfaces previously activated with UV/O_3_ (Figure 1A-1C). Uncoated regions were blocked by incubating the substrates for 5 min in a 1% Pluronic F-127 solution (BASF, Mount Olive, NJ) After blocking, the microprinted PDMS substrates were washed three times with a sterile PBS solution and stored at 4°C in PBS until use.

### 2.3 Neuronal networks of controlled architectures

Primary rat cortical neurons (RCN) prepared from the cortex of day-18 rat embryos (A10840-01, Life Technologies, Gaithersburg, MD) were suspended in a culture medium (21103-049, Neurobasal Medium, Thermofisher-Life Science) supplemented with 10 ml of B-27 supplement (17504044 + Thermofisher-Life Science), 1% antibiotic (15140-122, Life Technologies), 5 ml GlutaMax-I (35050-061,Thermofisher) and 50 μl of nerve growth factor (0.01 μg/ml, 556-NG, RD systems). Each culture corresponded to approximately 1.3 million cortical neurons obtained from day-18 rat embryos. RCN were seeded on microprinted substrates at a density of 30,000 cells/cm² and incubated under standard conditions at 37°celsius and 5 % CO_2_ (Figure 1D). The culture media was replaced the day after the seeding and then half-replaced every 48 hours until experiments were completed. Experiments were performed 6 hours after seeding (DIV0) and at days 4 (DIV4), 7 (DIV7), 12 (DIV12) or 13 (DIV13) post seeding.

### 2.4 Culture of primary cortical astrocytes

Rat cortical astrocytes were prepared from day-1 newborn rats. After the anaesthesia of the newborn, the brain was got back. Then the cortex was separated from the other parts of the brain and cut in pieces. To harvest cells, the pieces were gently mechanically dissociated by up down with a pipette and the cellular suspension was then filtered through a sieve and centrifuged 5 min at 1200 rpm to be washed. In order to purify the cells, the base was suspended in PBS-glucose (3 ml) and 1.5 ml were gently transfer on a Percoll gradient (Percoll 60%-Percoll 30%, Sigma Aldrich) and centrifuged 20 min at 1700 rpm. Cells were then getting back at the surface of the Percoll 30 % and suspended in PBS-glucose to be centrifuged at 1700 rpm during 20 min to remove the Percoll. At the end the cells were resuspended in 2 ml of medium (DMEM GlutaMax supplemented by 10 % FBS and 1 % antibody from Thermo Fisher Scientific, Invitrogen), counted via the trypan blue method and put in T-75 flask with approximately 20 ml of medium. The medium is fully replaced on the day after and then half-replaced two times a week (Fig. 2A).

**Figure 2.**
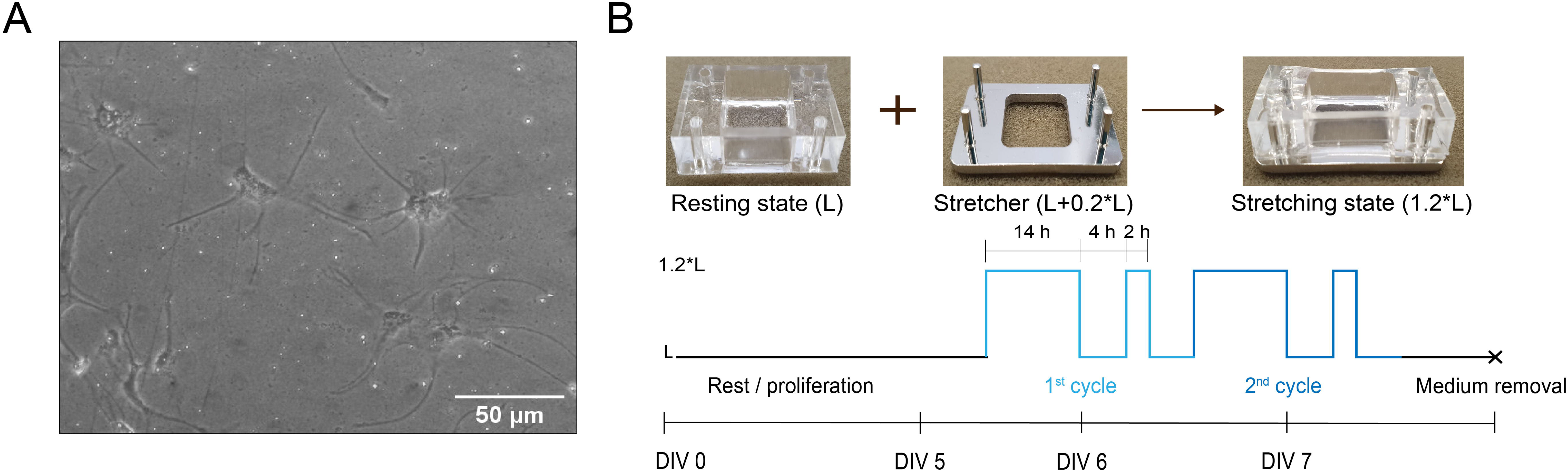
Primary cortical astrocytes were mechanically injured by stretching. (A) Primary astrocytes were cultured on thon PDMS membranes attached on a stretch chamber and homogeneously coated with PLL-LA. (B) Stretch chambers in resting state (L) were stretched at 20% to the stretching state (1.2 L) for cycle of 24 hours composed of 14 hours stretch, 4 hours rest, 2 hours stretch and 4 hours rest. Culture media were collected on DIV 7.

### 2.5 *In vitro* mechanical activation of astrocytes

To reproduce mechanical injuries, stretchable devices were made with PDMS moulded over a silanized Teflon mold. PDMS membrane of 150 μm thick was attached to the moulded pieces (Figure 2B). The membrane was realised by PDMS spin coating onto a silicon wafer (Fig. 1A). The field deformation of the elastic membranes was estimated by printing fluorescent protein (FITC-BSA) circles of 2000 μm² on the membrane of the device. The device was then submitted to a 20 % stretch along the horizontal axis and the distances between the centres of the circles were determined along the horizontal and the vertical axis. Our results report an elongation of ~20.4% along the horizontal axis and a slight shrinking of ~3.5% along the perpendicular axis of the stretched membrane (Supplementary Fig. S1).

After reaching the confluency in a T-75 flask, astrocytes were seeded on laminin/poly-L-lysine coated PDMS membrane at a concentration of 40,000 cells/cm² and the growing medium containing 3% of serum was added. On the fifth day, the device was stretched according to a cycle that consist of 4 successive steps: (i) one night of stretch, (ii) two hours of rest, (iii) four hours of stretch and (iv) two hours of rest. Mechanically activated astrocytes were submitted to either one (low activated) or two cycles (high activated) of stretch (Fig. 2B).

### 2.6 Cytokine quantification

A MSD multispot assay (Meso Scale Diagnostics, Maryland, US) was performed to quantify the amount of inflammatory cytokines and chemokines released by activated and control astrocytes. The MSD kit permitted to quantify the following cytokines: interferon gamma (IFN-γ); interleukins 4, 5, 6, 10, 13 and 1β (IL-4, IL-5, IL-6, Il-10, IL-13 and IL-1β); tumour necrosis factor alpha (TNF-α) and the chemokine KC/GRO also known as CXCL1, even at a very low concentration (lowest LLOD is 0.65 pg/ml for the IFN-γ). The used multi-array technology combines electrochemiluminescence and multi-spot plates to enable precise quantitation of multiple analytes in a single sample requiring less time and effort than other assay platforms. The assay can be considered as a "sandwich immunoassay" with a 96-well 10 spot-plate pre-coated with capture antibodies. The entire protocol is presented in Fig. S2.

### 2.7 Treatment of developing neuronal networks with cytokines

At the end of the second stretch cycle, the membrane was allowed to rest for four hours. Depending of the experiments to be performed, the supernatant of stretched astrocytes (one cycle or two cycle stretch) was added to neuronal networks at DIV7. Treated neuronal networks were let to grow until DIV13 then, the targeted immunostaining was performed. Immunostaining and cell viability assays of the astrocytes was performed directly after the stretch cycle(s). Neuronal networks were also incubated with only TNF-α at the concentration of 50 pg/ml, which represents the average amount of cytokine released by mechanically activated astrocytes. The TNF-α was added to the neuronal medium in two ways: with 2 doses (added at DIV7 and 11) or 4 doses (added at DIV4, 7, 9 and 11). Immunostaining experiments were performed at DIV13 for both protocols.

### 2.8 Immunostaining of cortical neurons and astrocytes

For neuron immunostaining, cells were fixed with paraformaldehyde 4 % in PBS for 10 min and permeabilized with 0.02 % Triton X100 in PBS for 10 min. Cells were then incubated with a blocking solution containing 5 % BSA in PBS. After three washes, cells were labelled with anti-synapsin Oyster 550 conjugated for staining synapsin (1:600, 106 011C3, Synaptic Systems), anti-βIII-tubulin Alexa Fluor 488 conjugated for staining microtubules (1:300, AB15708A4, Millipore) and DAPI for staining nuclei for one night at 4°C. For labelling actin and glial fibrillary acidic protein (GFAP), astrocytes were fixed with paraformaldehyde 4 % in PBS during 15 min and permeabilized with 0.5% Triton X100 5 min at room temperature. Cells were then incubated in a foetal bovine serum solution (FBS,5%) as blocking solution for 30 min at RT. After three washes, astrocytes were labelled 1 hour at RT with anti-GFAP (1:150,644702, BioLegend) for staining GFAP, phalloidin conjugated with Alexa Fluor 488 to stain F-actin and DAPI to visualize the nuclei. A second incubation with a secondary antibody tetramethylrhodamine-labelled was performed during 1h at room temperature. For labelling the two TNF alpha receptors, TNFR1 and TNFR2, neurons were fixed with paraformaldehyde 4% supplemented with 4 % sucrose in PBS during 15 min at 4°C. Cells were then permeabilized and blocked with 5% BSA supplemented with 0.5 % Triton X100 during 30 min at RT. After three washes, cells were labelled with anti-TNFR1 and anti-TNFR2 (1:400, sc-8436, sc-7862, Santa Cruz) during one night at 4°C. A second incubation of 1 hour at room temperature was then performed with goat anti mouse Alexa Fluor 555 and anti-rabbit FITC conjugated (1:400, F7512, Sigma Aldrich). For all the labelling, coverslips were finally mounted on microscope glass slides with Slow Fade Gold Antifade (Molecular Probes, Invitrogen).

### 2.9 Epifluorescence and confocal imaging

Immunostained preparations were observed in epifluorescence and confocal mode with an inverted Nikon Eclipse Ti-E motorized microscope (Nikon, Japan) equipped with a Nikon C1 scanhead, x10 Plan Apo (NA 1.45), x40 Plan Apo (NA 1.45, oil immersion) and x60 Plan Apo (NA 1.45, oil immersion) objectives, two lasers (Ar-ion 488 nm; HeNe, 543 nm) and a modulable diode (408 nm). Epifluorescence images were captured with a Roper QuantEM:512SC EMCCD camera (Photometrics, Tucson, AZ) using NIS Elements AR (Nikon, Japan).

### 2.10 Image analysis

All images were taken with NIS-Elements Advanced Research software (v.4.5, Nikon, Japan) using similar illumination and recording conditions (camera frequency, gain and lamp intensity). The quantification of the ratio of synapsin to tubulin area was performed in NIS Elements Advanced Research (v4.5, Nikon, Japan) and Matlab with different 5 areas obtained from 3 cultures (n=15). For neuronal networks treated with astrocytes media and TNF-α, we used 3 different cultures with n=25 (TNF-α), n=40 (+control astrocytes media) and n=24 (+activated astrocytes media).

For the corrected total fluorescence intensity of the GFAP fibres, images were taken with NIS-Elements Advanced Research software (v.4.5, Nikon, Japan) using similar illumination and recording conditions (camera frequency, gain, and lamp intensity) and a x40 Plan Apo objective. The area, the raw integrated density, the mean grey value and the number of cells were measured for each image. In addition, five random background regions were selected to obtain a mean grey value of the fluorescent background. The corrected total fluorescence was calculated using the following equation:

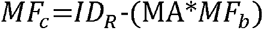

with *MF*_*C*_ was the corrected micropattern fluorescence, *ID*_*R*_ the raw integrated density, MA the micropattern area and *MF*_*b*_ the mean fluorescence background. This method was also applied to determine the fluorescence intensity of TNFR1 and TNFR2 receptors using a x60 Plan Apo objective.

### 2.11 Statistical analysis

Experimental data were presented using a boxplot representation and statistical comparisons were realized by a Student Test, Anova or via a Welsh test when the samples population number or the SD’s were significantly different between the two populations with *p < 0.05, 0.01 ≤ **p ≤ 0.05, ***p < 0.01.

## 3 Results

### 3.1 Glial fibrillary acidic protein as a marker of inflammation

Astrocytes contain a dense network of intermediate filaments, such as the glial acidic fibrillary protein (GFAP). A large number of studies have reported the upregulation of these intermediate filaments as an astroglial response to the injury. Impairment in the upregulation of these filaments have been shown to attenuate reactive gliosis and impaired formation of glial scar suggesting that the GFAP upregulation is an important parameter of the astrogliosis. The mechanism leading to this upregulation is still poorly understood but one explanation could be the elevation of the internal cell stiffness (D’arcangelo et al. 2000; Mochida 2015; Gruol, Vo, and Bray 2014). We stained for actin, GFAP and DAPI at DIV7 control cultures and activated cultures that underwent 1 and 2 cycles of stretch. We observed that most of the activated cells were positive (Fig. 3B) compared to control (Fig. 3A), suggesting that a mechanical stretch can activate astrocyte cultures. As shown in Fig. 3C, the GFAP area increased significantly after two stretches (2254 ± 806 μm²), whereas one cycle of stretch lead to a GFAP area (1018 ± 417 μm²) similar to controls (1043 ± 167 μm²). Interestingly, the GFAP fluorescence intensity after 1 cycle of stretch (49842 ± 12088 a.u) was similar to the controls (56848 ± 41974 a.u.), whereas 2 cycles of stretch astrocytes lead to a large increase of GFAP intensity (121754 ± 36923 a.u.) (Fig. 3D). Altogether, these results demonstrated that the production of GFAP is mechanosensitive and can be used as a marker of inflammation in astrocytes.

**Figure 3.**
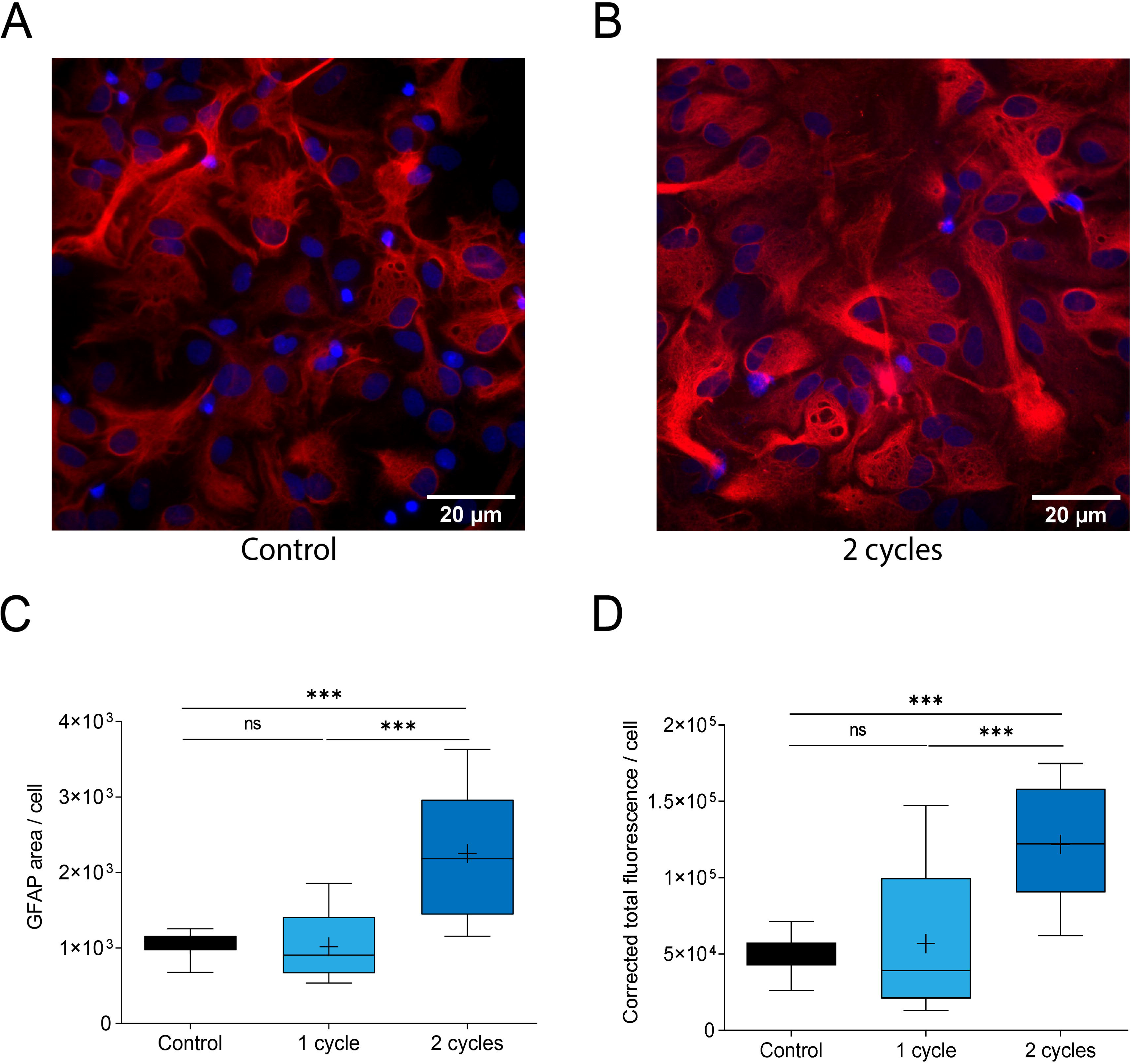
Mechanical activation of primary cortical astrocytes. Fluorescent images of GFAP and DAPI immunostained (A) control astrocytes and (B) 2 cycle stretched astrocytes. (C) GFAP area (μm²) per cell for control astrocytes (n=14), 1 cycle stretched astrocytes (n=24) and 2 cycle stretched astrocytes (n=19). (D) Corrected total fluorescence intensity (a.u.) per cell. Unpaired t-test, ns > 0.05, ***p < 0.001.

### 3.2 Quantification of the cytokines released by mechanically activated astrocytes

We analyzed the concentration of cytokines in control astrocytes and cultures that underwent 1 or 2 cycles of stretch (Fig. 4A). Our results indicated that control and activated astrocytes released five main cytokines and chemokines (IL-5, IL-6, IL-1β, TNF-α and KC/GRO) the others being negligible (C≤1 pg/ml). The concentrations of the inflammatory molecules were significantly lower for control than for astrocytes stretched two times, with 19.53 ± 8.9 and 68.3 ± 31.3 pg/ml for TNF-α (Fig. 4B), 7.57 ± 1.52 and 28.5 ± 7.5 pg/ml for IL-1β (Fig. 4C), 7.55 ± 0.8 and 11.05 ± 2 pg/ml for IL-5 (Fig. 4D), 89.2 ± 1.7 and 642.2 ± 320 pg/ml for IL-6 (Fig. 4E) and 1036 ± 94.2 and 4288 ± 258 pg/ml for KC/GRO (Fig. 4F). Interestingly, activated astrocytes released a much higher concentration of proinflammatory molecules that control cultures, suggesting that a mechanical stretch can induced a reactive state of the astrocytes, as observed in TBI. Our results show that the concentration of TNF-α, IL-1β, IL-5 and KC/GRO was not different for 1 cycle of stretch. Surprisingly, we observed that the concentration of IL-6 increased significantly after 1 cycle of stretch, suggesting that the expression of IL-6 is mechanosensitive. This hypothesis was confirmed by performing 2 cycles of stretch that lead to significant increase of IL-6. In addition, we observed a large increase of the concentrations of TNF-α, IL-1β and KC/GRO after 2 cycles of stretch, with a four time increase for TNF-α and KC/GRO. One may note that IL-5 exhibited the smallest increase of concentration after 1 and 2 cycles of stretch. In the following, we will focus our attention on the role of TNF-α, which is considered as an important marker of severe TBI (Adrian et al. 2016; Liu et al. 2014; Oshima et al. 2009).

**Figure 4.**
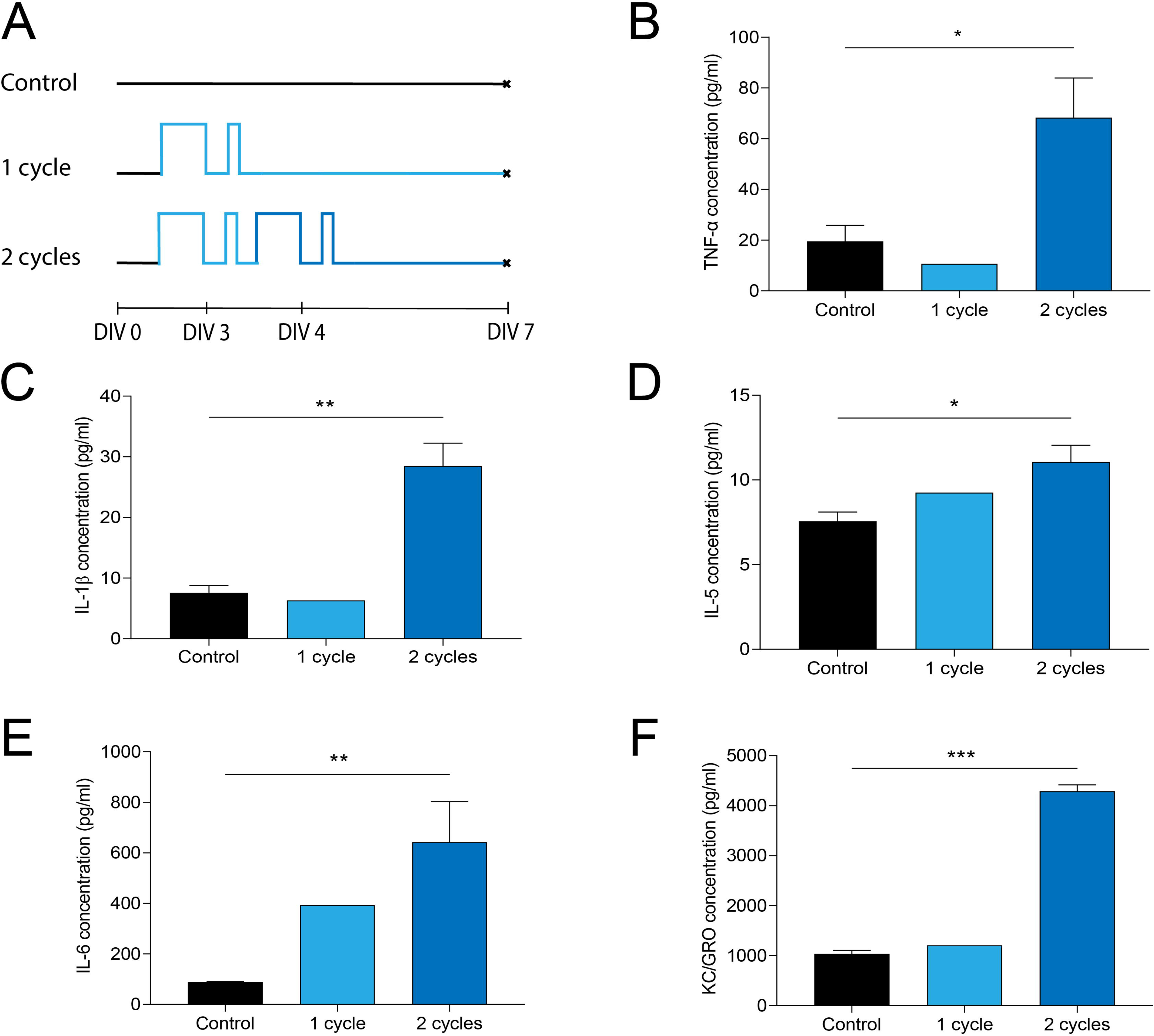
Determination of the chemokines and cytokines secreted by mechanically injured astrocytes. (A) Schematic timeline of experiments on control astrocytes, 1 cycle stretched astrocytes and 2 cycle stretched astrocytes. MSD multispot assay results for (B) TNF-α, (C) IL-1β, (D) IL-5, (E) IL-6 and (F) KC/GRO in control astrocyte medium (n=3), 1 cycle astrocyte medium (n=1) and 2 cycles astrocyte medium (n=2). The concentration of other cytokines was negligible (C≤1 pg/ml). Unpaired t-test, 0.01 ≤ *p ≤ 0.05, 0.001 ≤ **p ≤ 0.01, ***p < 0.001.

### 3.3 Effects of neuromodulatory cytokines on developing neuronal networks

The medium of control and activated astrocytes was added to cortical neuronal networks at DIV7 (Fig. 5A-B). Then neuronal networks were let to grow and immunostaining experiments were performed at DIV13. We performed Live/Dead assays on control neuronal networks and neuronal networks cultivated either with control astrocytes media (Fig. 5C) or activated astrocyte media (Fig. 5D). Our results indicated that control and activated astrocytes media did affect the neuron viability (Fig. S3B). Neuronal networks were stained at DIV13 for β-III tubulin and synapsin, which was used as a marker of presynaptic terminals. Figure 5E showed an increase of the ratio between β-III tubulin and synapsin area for neuronal networks cultivated with the astrocyte media obtained after two cycles of stretch (1.04 ± 0.62) in comparison with control neuronal networks (0.66 ± 0.21) and neuronal networks exposed to control astrocyte medium (0.67 ± 0.27). Our findings suggested that chemokines and cytokines secreted by astrocytes after two cycles of stretch can have a beneficial effect on neuronal networks, as observed previously (Figiel, 2008; Gougeon, 2013).

**Figure 5.**
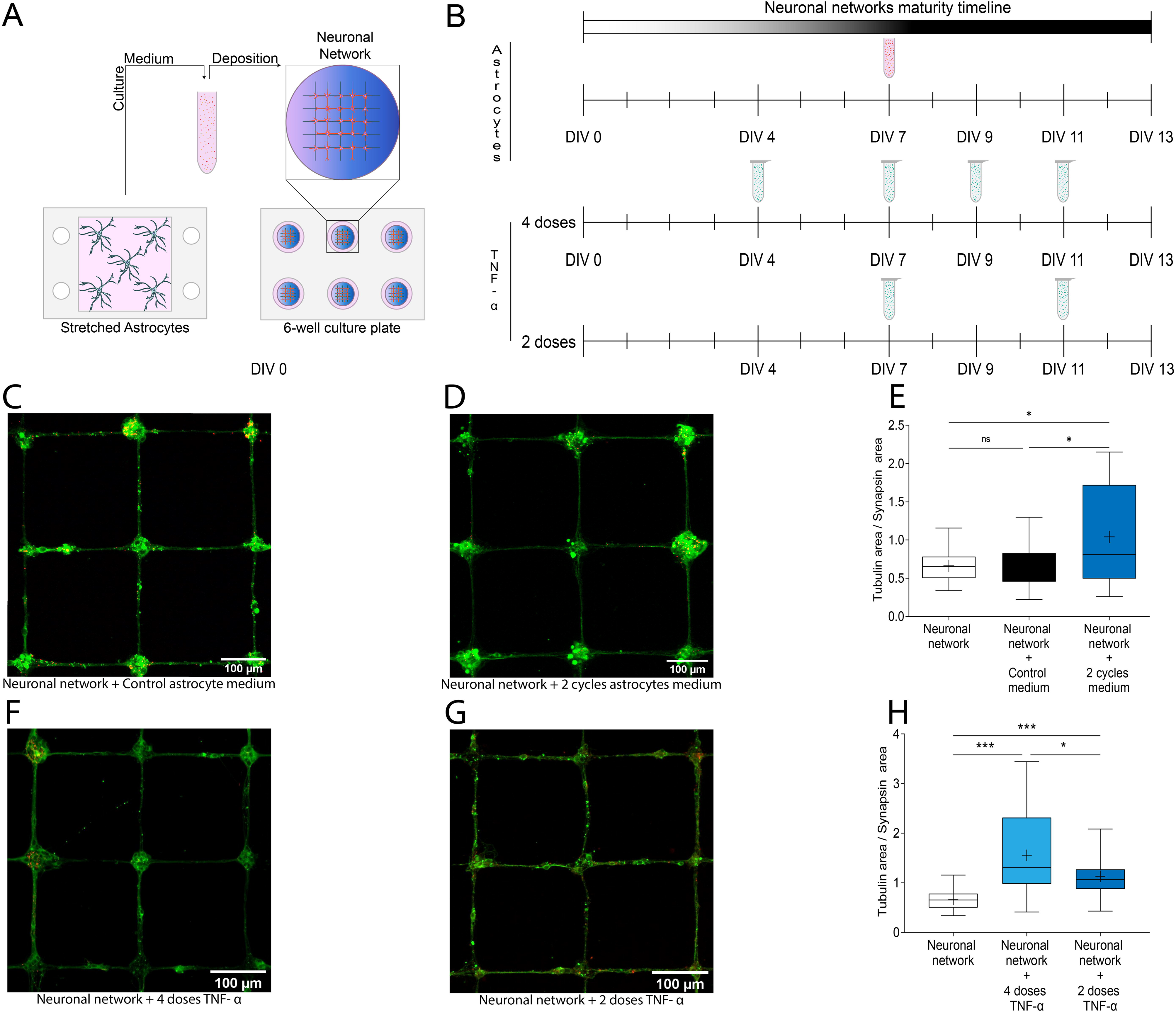
Neuronal networks were treated with the media of injured astrocytes and TNF-α. (A) The medium of stretched astrocytes was transferred to neuronal networks in 50/50 dilution. (B) Modulation of neuroinflammation was triggered either by stretched astrocytes medium or by 2 or 4 doses of TNF-α at regular time points. Fluorescent images of neuronal networks treated (C) with control astrocyte medium and (D) 2 cycle stretched astrocyte medium, both were immunostained for tubulin (in green and synaptophysin (in red). (E) Tubulin and synapsin ratio for control neuronal networks (n=16), neuronal networks with control astrocyte medium (n=41) and neuronal networks with 2 cycle stretched astrocyte medium (n=23). Fluorescent images of neuronal networks treated with (F) 4 doses of TNF-α and (G) 2 doses of TNF-α, both were immunostained for tubulin (in green) and synaptophysin (in red). (H) Tubulin and synapsin ratio for control neuronal networks (n=16), neuronal networks with control astrocyte medium (n=24) and neuronal networks with 2 cycle stretched astrocyte medium (n=26). Unpaired t-test, ns > 0.05, 0.01 ≤ *p ≤ 0.05, ***p < 0.001.

In order to decipher the individual contribution of TNF-α, we cultivated neuronal networks during 13 days with the addition of 4 doses of TNF-α added at DIV4, DIV7, DIV9 and DIV11 (Fig. 5F) or 2 doses of TNF-α added at DIV 7 and DIV11 (Fig. 5G). Live/Dead assays performed on healthy neuronal networks and on networks treated with 4 doses of TNF-α indicated that the neuronal cell viability was not affected (Supplementary Fig. S2). As shown in Fig. 5H, we found that the ratio between the tubulin area and the synapsin area was significantly higher in neuronal networks treated with either 4 (1.557 ± 0.814) or 2 doses of TNFα (1.132 ± 0.385). Our findings suggested that TNF-α can promote the tubulin polymerization by the stabilization of the microtubules (MTs). This effect of TNF-α on tubulin polymerization is supported by recent reports about the TNF-α mediated phosphorylation of the oncoprotein 18 (Vancompernolle, 2000). This pathway is probably not the only one conducting to the MT stabilization as TNF-α might also modulate other MAP proteins. The NFκB signalling pathways could also have an impact on the tubulin area and neurite outgrowth, as several studies reported evidences that the nuclear factor plays a role in regulating neurite outgrowth (Sole, 2004).

### 3.4 Modulation of the neuronal TNF receptors by injured astrocytes

Our results indicated that TNF-α plays an important role in the inflammatory response induced in neurons. This cytokine binds to two receptors, TNFR1 and TNFR2, leading to different signalling pathways: (i) the cell survival via the activation of the Nuclear Transcription Factor (NFκB) and (ii) the cell apoptosis via the caspase signalling. TNFR1 is expressed in almost all cell types and can promote cell survival or induce cell apoptosis upon the linkage with TNF-α. TNFR2 is mainly expressed in neurons and glial cells and promotes only cell survival (Naudé, 2011).

An important question that needed to be addressed was therefore to understand the individual role of TNFR1 and TNFR2 receptors in the modulation of the neuronal networks by injured astrocytes and TNF-α. To address this issue, we first investigated the effect of control and activated-astrocyte media on the balance between TNFR1 and TNFR2 (Fig. 6A). Based on immunostained images of each receptor, we determined the corrected total fluorescence of TNFR1 and TNFR2. Control neuronal networks were immunostained at DIV4, DIV7 and DIV12, whereas neuronal networks treated with either the astrocyte medium obtained from the control and two cycle stretched astrocytes were stained at DIV12. First of all, we observed that the amount of R1 and R2 receptors increased with the maturation of neuronal networks from DIV4 (40780 ± 14664 a.u) to DIV12 (1271480 ± 40869 a.u), while keeping a perfect balance between both receptors at DIV12 (Fig. 6A). The addition of medium obtained from control astrocytes did not statistically disturb the balance between TNFR1 and TNFR2, suggesting that healthy astrocytes did not affect both receptors. However, the addition of astrocyte medium obtained from a two cycle stretch assay exhibited a significant modulation of the amount of TNFR1 and TNFR2, leading to a large imbalance between both receptors. Indeed, we found that astrocyte medium obtained from a two cycle stretch assay decreased the TNFR1 fluorescence signal from 71482,1 ± 34699,5 a.u. for control at DIV12 to 52440,4 ± 28481,3 a.u. whereas we observed a significant increase of the TNFR2 fluorescence signal from 74438,5 ± 30995,7 a.u. for control at DIV12 to 103122 ± 53886 a.u. (Fig. 6A). Altogether, our findings indicated that cytokines and chemokines expressed by injured astrocytes lead to a significant modulation of TNFR1 and TNFR2 receptors, characterized by a decrease of TNFR1, that can promote cell survival or can induce cell apoptosis, and a large increase of TNFR2 that promotes only cell survival.

**Figure 6.**
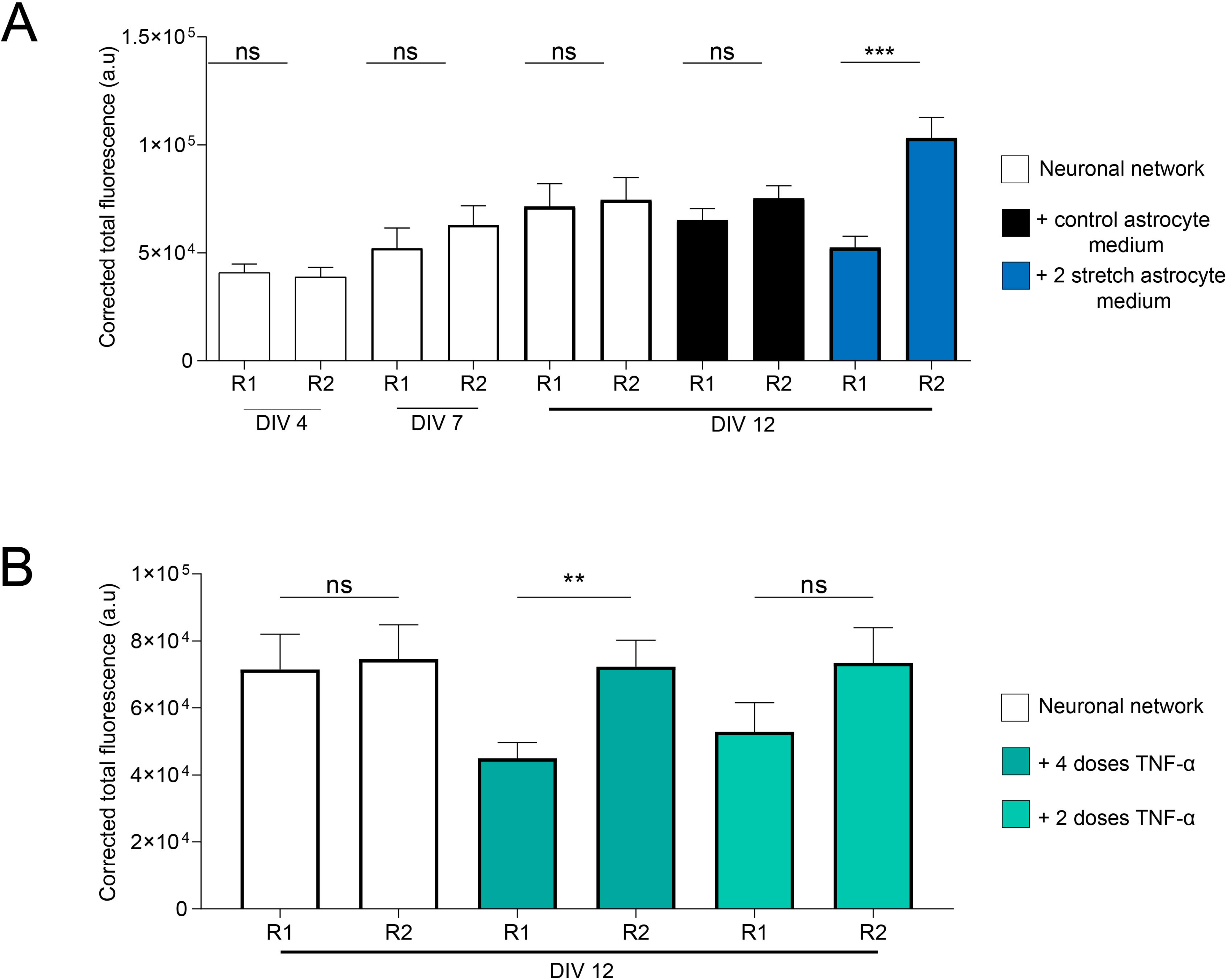
Determination of the amount of R1 and R2 TNF-α receptors in neuronal networks modulated by the media of mechanically injured astrocytes or doses of TNF-α. (A) Corrected total fluorescence of R1 and R2 TNF-α receptors in neuronal networks at DIV4 (n=13), DIV7 (n=14) and DIV12 (n=15). The corrected total fluorescence of TNF-α receptors for neuronal networks grown with control astrocyte medium (n=68) or 2 cycle astrocyte medium (n=43) was calculated at DIV12. One-way ANOVA (=5.218 and r² = 0.1381). (B) Corrected total fluorescence of the TNFα receptors for control neuronal networks and networks treated with either 4 (n=26) or 2 doses of TNFα (n=19). The corrected total fluorescence is presented at DIV 12. One-way ANOVA (F=2.83 and r² = 0.1354). ns > 0.05, 0.01 ≤ *p ≤ 0.05, 0.001 ≤ **p ≤ 0.01, ***p < 0.001.

Based on this observation, we further investigate the individual role of TNF-α in the antagonist modulation of TNFR1 and TNFR2 receptors. Interestingly, we found that the perfect balance between R1 and R2 receptors at DIV12 was statistically significantly affected by the addition of 4 doses and 2 doses of TNF-α. Indeed, the addition of TNF-α significantly decreased the amount of TNFR1 from 71482 ± 40869 a.u. to 44966 ± 19993 a.u. at DIV12 with 4 doses and to 52877 ± 31742 a.u. with 2 doses. Our results demonstrated that the addition of TNF-α decreased the expression of TNFR1 but did not affect the expression of TNFR2 that promotes cell survival. Altogether, our results suggested that the modulation of the neuronal networks by TNF-α expressed in injured astrocytes is based on a lower expression of TNFR1 receptors.

## 4 Discussion

This study shows a novel approach for studying mechanical-dependant release of inflammatory molecules by astrocytes by combining stretching assays and neuronal networks of controlled architectures. Rat cortical astrocytes were cultivated on stretchable devices for 5 days and submitted to a 20 % stretch following a stretching cycle for 2 days. Cortical neuronal networks were then grown with astrocyte media obtained from control and injured cultures. We have shown that a two cycle stretch induces a reactive phenotype of astrocytes, as observed in traumatic brain injuries (Burda, 2016). Our results showed that GFAP can be used as a marker of inflammation in injured astrocytes and suggest that a single stretch does not trigger the activation of astrocytes. In this context, it would be then valuable to consider in the future the role of microglia in the activation of astrocytes.

We demonstrated that stretch cycles triggered the release of proinflammatory cytokines such as IL-1,5,6 and −β, TNF-α and chemokines such as KC/GRO. Taken together, our results suggest that stretching assays on astrocyte cultures represent a valuable way to reproduce inflammatory conditions. While many reports presented TNF-α and IL-6 as neurotrophic agents (Figiel, 2008; Hirota, 1996), other described the negative effects of TNF-α on neurite outgrowth and branching of hippocampal neurons (Naudé, 2011; Bradley, 2008). In this context, our findings report that the cytokines and chemokines released by the reactive astrocytes regulate tubulin and synapsin areas to provide a beneficial effect on neuronal networks.

We investigated the effect of the media obtained from injured astrocyte cultures on TNFR1 and TNFR2 receptors. Our findings indicate that the amount of both receptors increases with the maturation of the network from DIV 4 to 12. In addition, we observed a significant modulation of the amount of TNFR1 and TNFR2 in response to the media obtained from injured astrocyte cultures, leading to a large imbalance between both receptors. In addition, we showed a modulation of the expression of the TNFR2 when neuronal networks were incubated with injured astrocyte medium, while TNFR2 remained constant in response to TNF-α. This may be due to the effect of the metalloprotease TACE, which is known to shed membrane-bound TNF-α but also the TNFR2 (MacEwan, 2002; Higuchi and Aggarwal, 1994). In this way, the cell is capable to determine its sensitivity to the extracellular TNF-α and so determine the outcome of the binding between the cytokine and its receptor.

Besides, the viability assays performed on our neuronal networks submitted to either astrocyte media or TNF-α showed no impact of the pro-inflammatory cytokines on the neuronal viability. Taking into account the positive impact of the cytokines on the growth of the neuronal networks and those concerning the TNF receptor modulation, we could easily say that the NFκB pathway has been triggered in the neurons via the released cytokines. Indeed, it has been demonstrated by Mettang et al. in 2018 that the activation of the NFκB pathway in neurons has a neuroprotective effect (Mettang, 2018). Interestingly, the activation of the same pathway in microglia mediates the inflammation (Tao, 2018). One potential therapeutic target could then be this pathway with its specific activation in neurons and its inhibition in microglia.

In conclusion, we showed here an interesting way to decipher the role of mechanically-injured astrocytes on neuronal networks. This model offers the possibility to apprehend and further study the molecular mechanisms of neuroinflammation in a context of traumatic brain injury. Further studies on co-cultured neuronal networks with astrocytes and microglia are needed in parallel with the pharmaceutical screening of therapeutic drugs in order to circumvent the detrimental effects of the pro-inflammatory cytokines and improves the beneficial ones by eventually playing on the NFκB signalling pathway both in neurons and glia.

## 6 Author Contributions

S.G. and L.R. designed the study. J.L. and A.P. performed experiments and analyzed data. J.L., A.P. and S.G. interpreted results. J.L., A.P., S.G. and L.R. wrote the manuscript and prepared figures. L.R., A.V, S.H. and L.B. discussed the results. All authors improved the manuscript.

## 8 Acknowledgments

We acknowledge and Dr. Raphaëlle Caillerez from The Alzheimer & Tauopathies Team of The Jean-Pierre Aubert Research Centre in Lille for her work and help with the quantification of cytokines.

