## Supplementary Information for "Role of TNF-α among inflammatory molecules secreted by injured astrocytes in the modulation of *in vitro* neuronal networks"

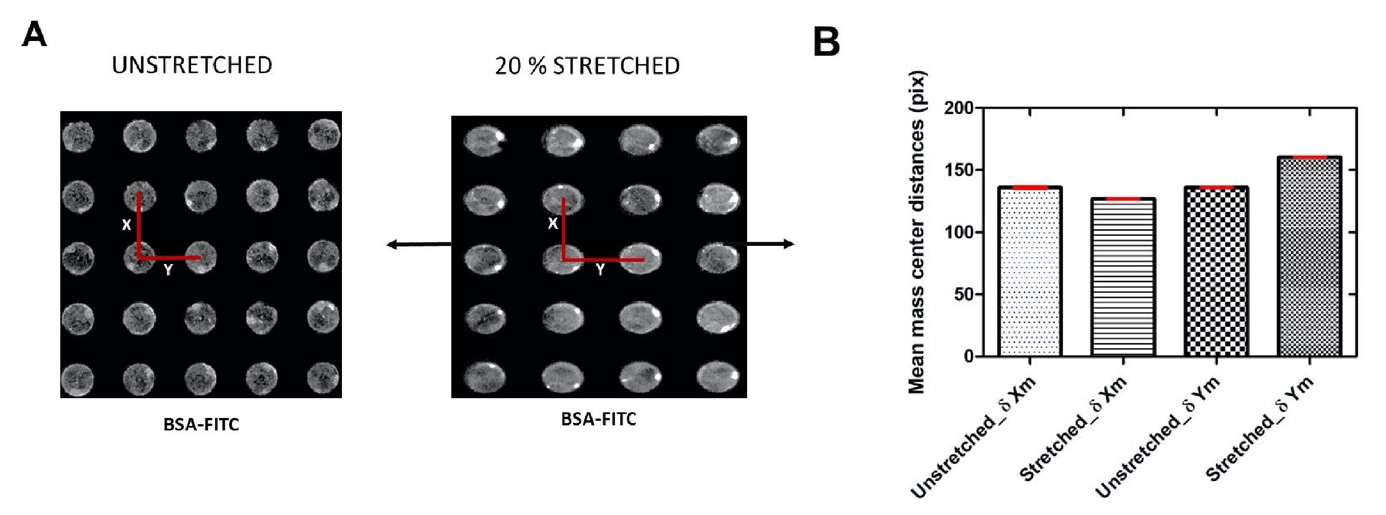

**Supplementary Figure 1 -** The field deformation of the elastic membranes was estimated by printing fluorescent protein (FITC-BSA) circles of 2000 µm² on the membrane of the device. (A) The device was then submitted to a 20 % stretch along the horizontal axis and the distances between the centres of the circles were determined along the horizontal and the vertical axis. (B) An elongation of ~20.4% along the horizontal axis and a slight shrinking of ~3.5% along the perpendicular axis of the stretched membrane was then determined from the mean mass center distances.

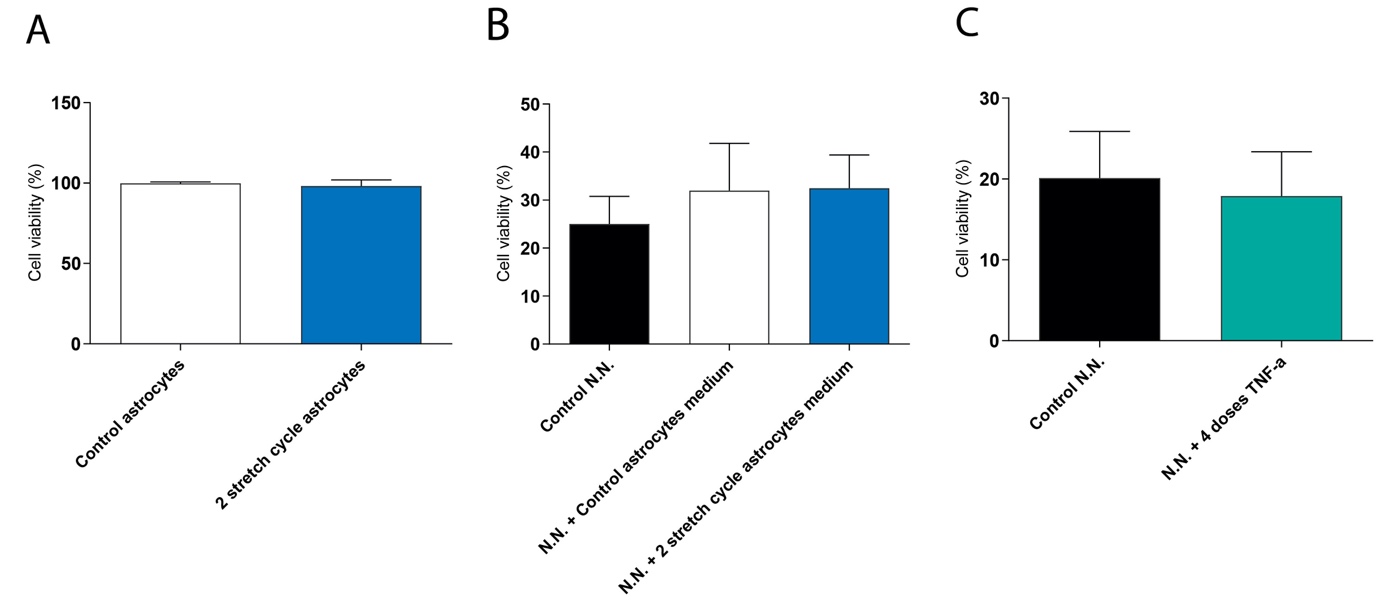

**Supplementary Figure 2 -** Live/Dead assays performed on (A) control astrocytes and 2 stretch cycle astrocytes, (B) control neuronal networks (N.N.), neuronal networks with control astrocyte medium and neuronal networks with 2 stretch cycle astrocyte medium and (C) control neuronal networks and neuronal networks treated with 4 doses of TNF-α.
